# A truncation mutant of adenomatous polyposis coli (APC) impairs apical cell extrusion through elevated epithelial tissue tension

**DOI:** 10.1101/2024.03.06.583801

**Authors:** Wan J. Gan, Rabina Giri, Jakob Begun, Helen E. Abud, Edna C. Hardeman, Peter W. Gunning, Alpha S. Yap, Ivar Noordstra

**Affiliations:** Centre for Cell Biology of Chronic Disease, Institute for Molecular Bioscience, The University of Queensland, St Lucia, Queensland, Australia; Mater Research - The University of Queensland, Woolloongabba, Queensland, Australia; Department of Gastroenterology, Mater Hospital, South Brisbane, Queensland, Australia; Department of Anatomy and Developmental Biology, Development and Stem Cells Program, Monash Biomedicine Discovery Institute, Monash University, Clayton, Victoria, Australia; School of Biomedical Sciences, Faculty of Medicine and Health, University of New South Wales, Sydney, New South Wales, Australia

## Abstract

Tissue tension encompasses the mechanical forces exerted on solid tissues within animal bodies, originating from various sources such as cellular contractility, interactions with neighbouring cells and the extracellular matrix. Emerging evidence indicates that an imbalance in tissue tension can influence structural organisation, homeostasis and potentially contribute to disease. For instance, heightened tissue tension can impede apical cell extrusion, leading to the retention of apoptotic or transformed cells. In this study, we investigate the potential role of adenomatous polyposis coli (APC) in modulating tissue tension. Our findings reveal that expression of an APC truncation mutant elevates epithelial tension via the RhoA/ROCK pathway. This elevation induces morphological alterations and hampers apoptotic cell extrusion in cultured epithelial cells and organoids, both of which could be mitigated by pharmacologically restoring the tissue tension. This raises the possibility that APC mutations may exert pathogenetic effects by altering tissue mechanics.

## INTRODUCTION

Solid tissues in animal bodies experience a variety of mechanical forces that can profoundly influence their form and function. This is well-understood for mechanical tension (Chan et al. 2019; Harn et al. 2019; Eder, Aegerter, and Basler 2017), defined as forces that tend to pull tissues apart (Charras and Yap 2018). Tissue mechanical tension guides cell behaviours such as proliferation, differentiation, and migration, thereby sculpting the mechanical characteristics and structural arrangement of tissues (Heer and Martin 2017; Le and Mayor 2023). Aberrant changes in tissue tension have been implicated in the pathogenesis of numerous diseases, including cancer, fibrosis, and cardiovascular disorders, highlighting its significance in disease progression and tissue pathology (Ayad, Kaushik, and Weaver 2019; Zuela-Sopilniak and Lammerding 2022). It is therefore important to understand the mechanisms that determine tissue tension and how these influence morphogenesis and tissue homeostasis.

The physiologic and pathologic impact of tissue tension has been extensively investigated in epithelia. Here a key determinant of tissue tension arises from interactions between the actomyosin cytoskeleton and E-cadherin adhesion at adherens junctions (AJ) (Campàs, Noordstra, and Yap 2023). The actomyosin contractile apparatus is the principal tension-generator of cells; its actin filaments physically bind to the classical cadherin adhesion system; and E-cadherin adhesions also regulate contractility by modulating cell signals, especially the RhoA GTPase (Ratheesh and Yap 2012; Yap, Duszyc, and Viasnoff 2018). Together, these cellular mechanisms allow actomyosin to generate mechanical tension at AJ in epithelial monolayers under steady-state conditions. As well, junctional contractility influences homeostatic events, such as the process of apical extrusion where cells are expelled from a tissue apically, as part of tissue homeostasis or defence mechanism to remove damaged cells. Apical extrusion is elicited by a range of homeostatic changes in an epithelium including apoptosis and expression of oncogenes. In both cases, extrusion is driven by changes in actomyosin activity at the junctions between apoptotic or oncogene-expressing cells and their immediate neighbours (Duszyc et al. 2021; Teo, Gomez, et al. 2020). Regulation of the AJ cytoskeleton then plays an important role in maintaining baseline tissue tension as well as the local changes necessary for apical extrusion.

Recently, we reported that the efficacy of apical extrusion is reduced when the pre-existing mechanical tension of epithelial monolayers is elevated. This applied for both apoptotic (Mann et al. 2024) and oncogenic extrusion (Teo, Gomez, et al. 2020), suggesting that cytoskeletal regulators of contractility can influence the nexus between baseline tissue tension and apical extrusion. Adenomatous polyposis coli (APC), a well-known oncogene which is best understood for its role as a regulator of the Wnt signaling pathway (Korinek et al. 1997; Morin et al. 1997; Gao et al. 2002; Flanagan et al. 2021; van Neerven et al. 2021), has been demonstrated to have the capacity to influence the cytoskeleton. It exerts such function through interactions with key regulators including mDia, EB1 and small GTPases, which play pivotal roles in modulating actomyosin contractility (Okada et al. 2010; Kawasaki et al. 2000; Juanes et al. 2020). Moreover, APC has been shown to participate in the regulation of AJ (Baro et al. 2023; Carothers et al. 2001). In this paper, we investigate how cytoskeletal regulation by the APC gene product can influence apical extrusion through the modulation of tissue tension.

## RESULTS AND DISCUSSION

### An APC truncation mutant increases epithelial monolayer tension

To investigate the role of APC in regulating epithelial monolayer tension, we utilized CRISPR-Cas9 technology to generate two distinct genetic modifications in MCF10A cells. Firstly, we created a complete APC knockout (APC^KO^) by introducing a frameshift at amino acid 17, resulting in a premature stop at amino acid 30 (Figure 1a, S1a-b). Secondly, we induced a frameshift mutation in exon 11, leading to the expression of a truncated form of APC (APC^Trunc^) (Figure 1a, S1a-b). This strategic alteration was guided by the recognition that a significant proportion of disease-associated mutations in APC are found within the mutation cluster region (MCR), often leading to the expression of N-terminal truncations of APC which contribute to pathological processes, such as gut polyp formation (Smith et al. 1993; Kawasaki et al. 2009; Yonemura et al. 2010; Nakamura 1995; Tominaga et al. 1998; Kohler et al. 2008).

**Figure 1:**
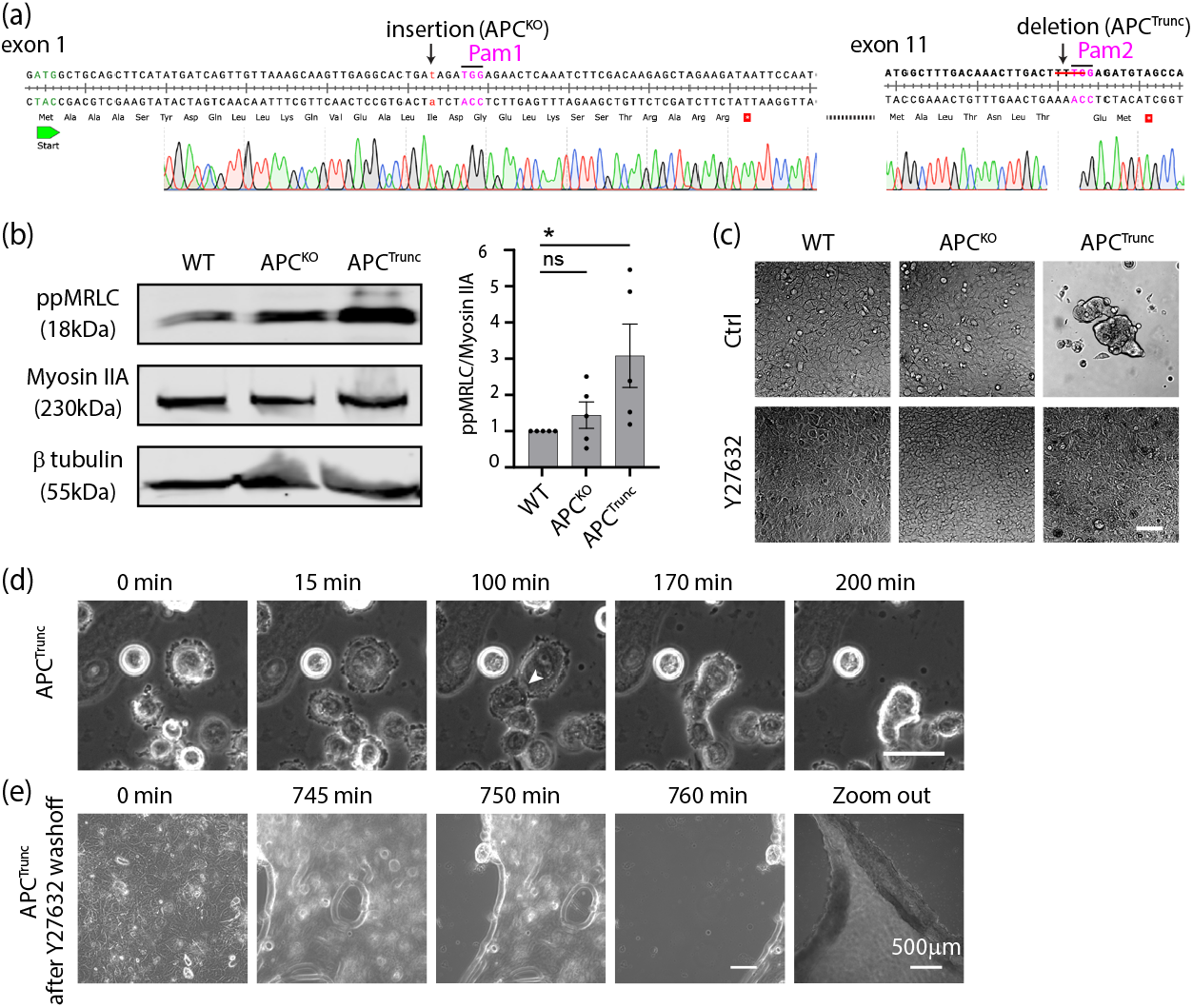
Truncation of APC increases epithelial monolayer tension. (a) Genomic sequencing of human *APC* exon 1 and exon 11 from MCF10A APC^KO^ and APC^Trunc^ cells respectively. (b) Western blot analysis and quantification of WT, APC^KO^ and APC^Trunc^ cell extracts detected for phosphorylated myosin light chain (ppMRLC), myosin IIA, and β tubulin. ^*^p=0.0346; ns:not significant; Kruskal-Wallis test. Data are means ± SEM, with individual data points indicated. (c) WT, APC^KO^, and APC^Trunc^ cells cultured on BME-II-coated PDMS substrates (10-20kPa). Cells were either untreated (Ctrl) or treated with 10µM Y27632. Scale bar: 100µm. (d) Live cell recording of APC^Trunc^ cells on BME-II-coated PDMS substrates (10-20kPa). Arrowhead indicates the initiation of cell-cell contact. Scale bar: 50µm. (e) Live cell recording of APC^Trunc^ monolayer after washing out the Y27632. Scale bar: 100µm. Scale bar of the zoom out image: 500µm.

Non-muscle Myosin II in the contractile apparatus is activated by phosphorylation of its regulatory light chain (MRLC) and generates force upon interaction with actin filaments (Krendel et al. 1999; Leerberg et al. 2014). To test if manipulation of APC affected actomyosin activity, we examined the phosphorylation status of MRLC by Western blot analysis. Cellular levels of ppMRLC, relative to Myosin IIA heavy-chain expression, were not altered in the APC^KO^ cells compared to wildtype (WT) controls. Strikingly, however, ppMRLC was elevated in the APC^Trunc^ cell line (Figure 1b), suggesting that expression of this N-terminal mutant could stimulate actomyosin. To characterize this further, we then examined how APC manipulation affected morphology of these cell lines.

Previous studies have illustrated that cells seeded on soft substrates undergo notable morphological changes and round up, particularly in response to heightened levels of tissue tension (Nyga et al. 2021). To investigate how increased tissue tension influences epithelial spreading on a soft substrate, we seeded cells onto BME-II-coated PDMS layers with a stiffness ranging between 10-20 kPa (Teo, Lim, et al. 2020). While WT and APC^KO^ cells efficiently spread to form monolayers on the substrate (Figure 1c, Movie 1a), APC^Trunc^ cells displayed a markedly different behaviour. Rather than spreading, these cells appeared to round-up to form 3D aggregates (Figure 1c, Movie 1b). Interestingly, detailed time-lapse imaging revealed that, after seeding, cells first spread but round up once the first cell-cell contacts are initiated (Figure 1d, arrowhead). To test if this morphological pattern was due to increased contractility in the cells, we used Y27632 to inhibit Rho kinase (ROCK), a principal activator of actomyosin contractility. ROCK inhibition reversed aggregation and rounding, allowing APC^Trunc^ cells to form a monolayer on the PDMS surface (Figure 1c, Movie 1c). However, this effect was transient as cells lost their substrate connections and reverted to their rounded morphology upon Y27632 washout (Figure 1e, Movie 1d).

Together, these findings suggest that expression of APC^Trunc^ might increase actomyosin contractility via the RhoA-ROCK pathway. To confirm this directly, we used pull-down assays to measure cellular levels of active GTP-RhoA. Indeed, GTP-RhoA levels were significantly increased in the APC^Trunc^ cells compared to WT control cells (Figure S1c). Overall, these findings indicate that the N-terminal APC^Trunc^ mutant enhances RhoA signaling to stimulate cell contractility.

### The APC N-terminus increases adherens junction tension

We then asked if the increased cell contractility was transmitted to AJ for mechanical tension. For this, we evaluated mechanosensitive conformational changes in α-catenin within the E-cadherin molecular complex. In the absence of mechanical tension, α-catenin adopts a closed conformation, which masks epitopes in its central M-domain and C-terminal actin-binding domain. Application of tension induces conformational changes that expose these epitopes, which can be detected by the α-18 mAb and VD7 pAb, respectively (Duong et al. 2021; Noordstra et al. 2023; Yonemura et al. 2010). Accordingly, we used these markers of molecular-level tension to test if AJ tension was altered by manipulation of APC. Cells were grown to confluence on glass coverslips to facilitate the formation of epithelial monolayers, then fixed and immunostained for α-catenin and its epitopes.

Interestingly, we observed intensified labelling with both α-18 and VD7 antibodies in APC^Trunc^ cells compared to WT and APC^KO^ cells (Figure 2a-b). This supports the notion that the increased tension is indeed reflected by a conformational change of α-catenin. To confirm that this mechanical change was due to the N-terminal fragment of APC, we exogenously reintroduced a GFP-tagged truncation of APC, similar to APC^Trunc^, in the APC^KO^ cells (Figure S1a, d). These cells were designated as APC^KO+N-APC^. Consistent with the increased junctional tension observed in APC^Trunc^, the APC^KO+N-APC^ monolayer exhibited an increase in α-18 and VD7 labelling as compared to the WT and APC^KO^ cells (Figure 2a-b).

**Figure 2:**
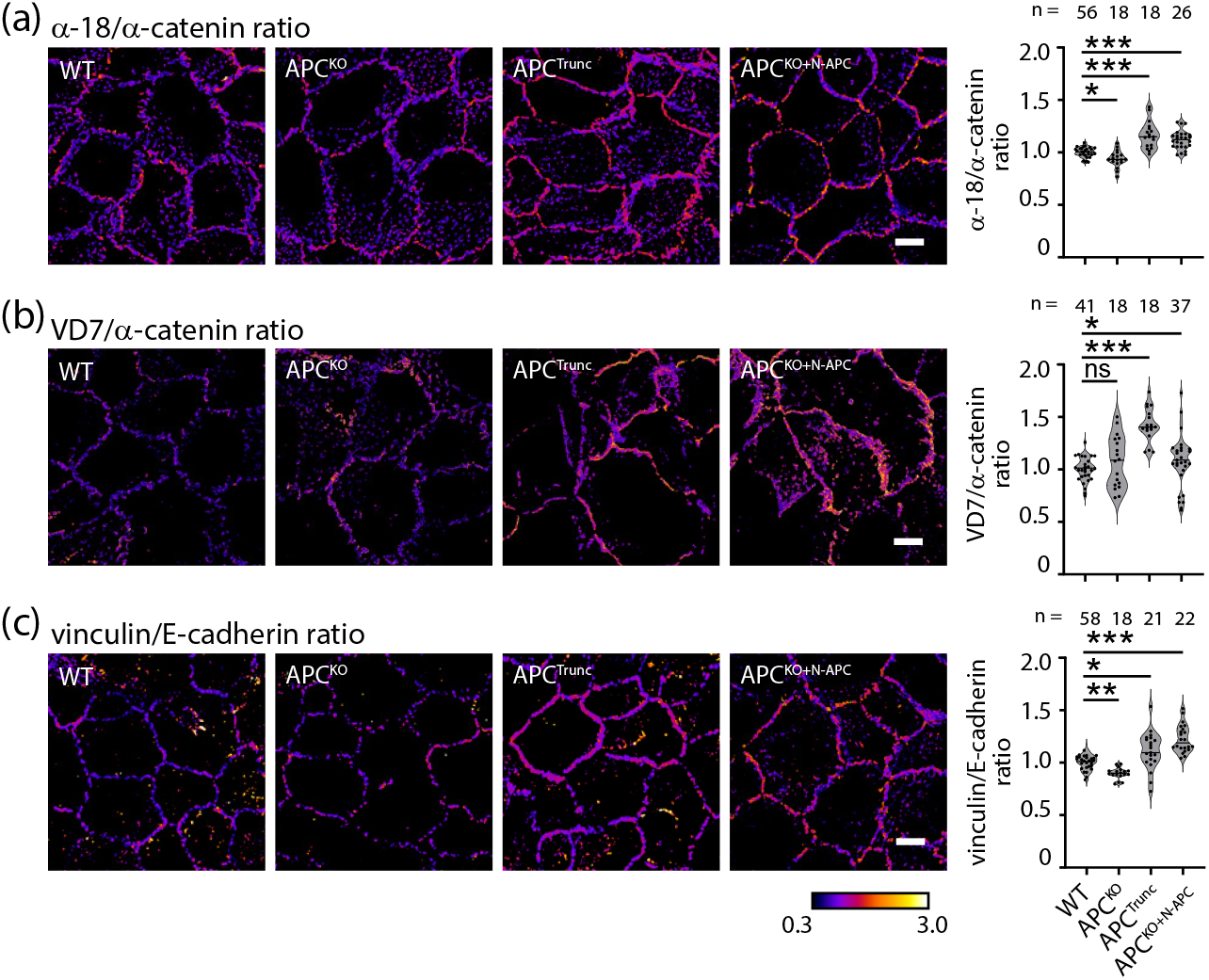
The APC N-terminus increases adherens junction tension. (a-c) Ratiometric images and quantifications of WT, APC^KO^, APC^Trunc^, and APC^KO+N-APC^ MCF10A cells stained for (a) α-18/α-catenin, (b) VD7/α-catenin, or (c) vinculin/E-cadherin respectively. Scale bar: 10µm. ^*^p<0.05; ^**^ p<0.01; ^***^: p<0.001; ns: not significant; Kruskal-Wallis test. Data are means ± SEM, with individual data points indicated, and are obtained from 3 independent experiments.

As a further test of tension at AJ, we immunostained for vinculin. Vinculin binds to the central domain of α-catenin when its conformation is opened on application of tension (Yonemura et al. 2010). Consistent with our previous findings, junctional vinculin labelling intensifies in APC^Trunc^ and APC^KO+N-APC^ cells compared to WT and APC^KO^ cells (Figure 2c). Altogether, our data indicate that baseline mechanical tension at AJ is increased when expression of N-terminal APC mutants stimulates contractility.

### The APC N-terminus compromises apical extrusion of apoptotic cells

We recently reported that tissue mechanical hypertension disrupted the ability of epithelial monolayers to eliminate apoptotic cells by apical extrusion (Mann et al. 2024). Therefore, we tested whether the tissue mechanical changes induced by APC^Trunc^ mutants also affected apoptotic extrusion.

We utilised E-cadherin tagged with mCherry to delineate individual cells within the monolayer. To induce apoptosis and monitor extrusion, we selectively injured the nuclei of single cells using a two-photon pulsed laser beam and subsequently imaged the cells for 1 hour following cell death. In WT monolayers, approximately 67% of the injured cells were extruded upon laser injury (Figure 3a-c, Movie 2a). The extrusion process was characterized by the apical expulsion of the apoptotic cell (identified with Annexin V) and the simultaneous extension of its neighbours to form a rosette that seals the epithelium (Figure 3a, b). APC^KO^ monolayers were able to efficiently extrude apoptotic cells with an apical extrusion rate that was greater than WT cells (Figure 3a-c, Movie 2b). This increase in extrusion efficacy may be due to altered adhesive properties, potentially making it easier for the cells to disengage from basal connections and be expelled from the epithelial layer.

**Figure 3:**
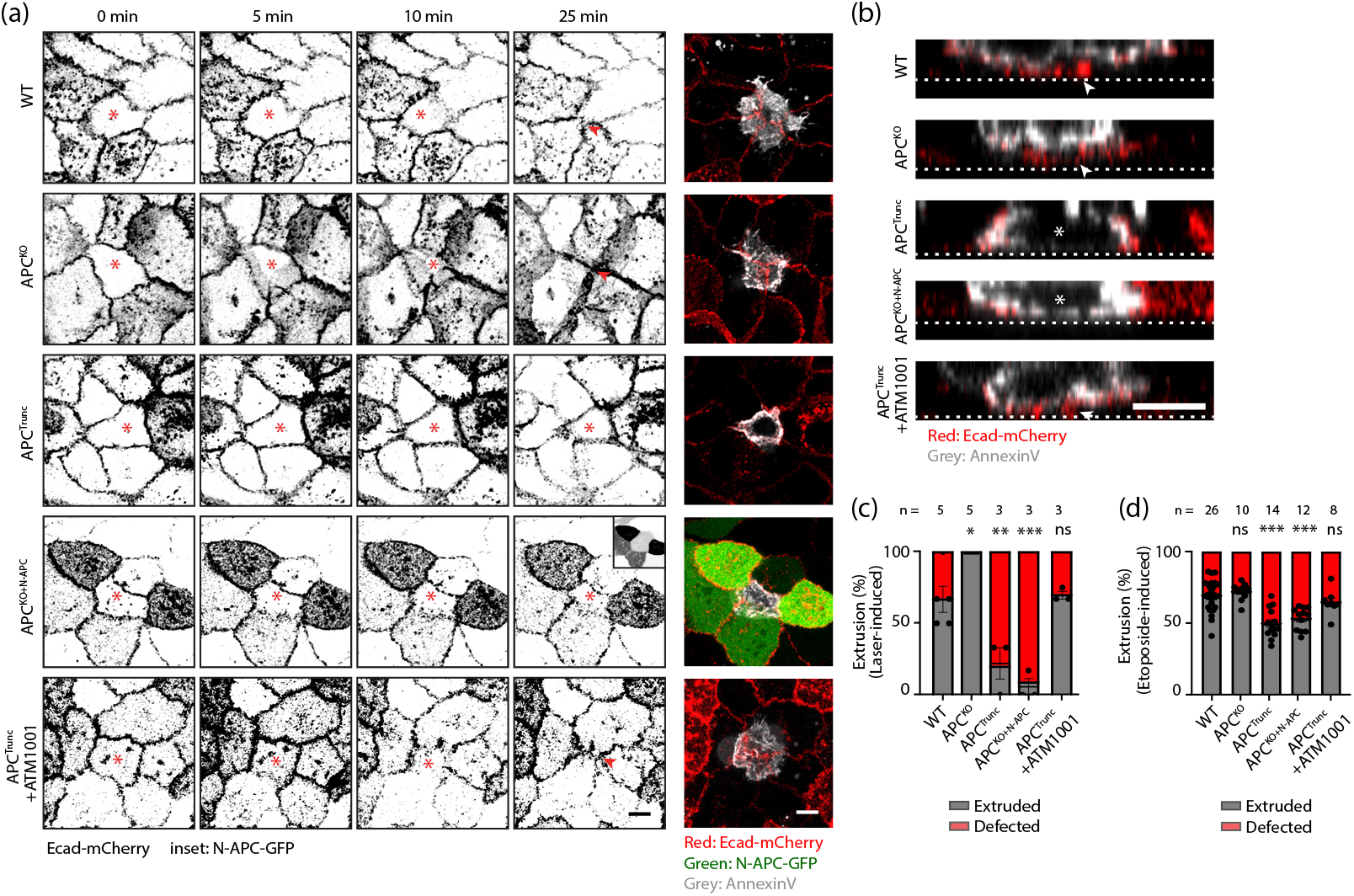
Expression of the APC N-terminus compromises apical extrusion of apoptotic cells. (a) Live cell recording of apoptotic cell extrusion within E-cadherin-mCherry (red) expressing WT, APC^KO^, APC^Trunc^, APC^KO+N-APC^ and ATM1001-treated APC^Trunc^ MCF10A cells. N-APC-GFP expression (green). Annexin V cell death marker (white). ^*^cell injured with two-photon pulsed laser; Red arrow: closure of the epithelial layer upon extrusion. Scale bar: 10µm. (b) Orthogonal views of cells in (a) at 25 min. *retained cells after laser injury; Red arrow: closure of the epithelial layer upon extrusion; white dotted line: glass surface. Scale bar: 10µm. (c) Quantification of apical cell extrusion as a result of laser injury as in (a) and (b). ^*^: p<0.05; ^**^: p<0.01; ^***^: p<0.001; ns: not significant; Kruskal-Wallis test. Data are means ± SEM, with individual data points indicated, and are obtained from 3-5 independent experiments. (d) Quantification of apical cell extrusion as a result of etoposide treatment. ^***^: p<0.001; ns: not significant; Kruskal-Wallis test. Data are means ± SEM, with individual data points indicated, and are obtained from 3 independent experiments.

In contrast, extrusion was markedly compromised in APC^Trunc^ cells, with extrusion rates falling below 20% (Figure 3a-c, Movie 2c). This was attributable to the APC truncation fragment, as the capacity of APC^KO^ monolayers to support apical extrusion was significantly compromised by expression of the APC N-terminal transgene (APC^KO+N-APC^, Figure 3a-c, Movie 2d). To test if aberrant tissue tension was responsible for this defect in apoptotic extrusion, we used the ATM1001 tropomyosin inhibitor, which reduces junctional contractility by disrupting the binding of tropomyosin 3.1 to actin filaments (Kee et al. 2018; Mann et al. 2024). Remarkably, a 24-hour treatment with 2.5µM ATM1001 effectively restored the apoptotic extrusion rate in the APC^Trunc^ monolayer to levels comparable to those observed in the WT cells (Figure 3a-c, Movie 2e).

As an alternative test, we investigated extrusion dynamics upon apoptosis induced with the podophyllotoxin etoposide (Karpinich et al. 2002). No difference was observed between WT and APC^KO^ cells. Nonetheless, APC^Trunc^ and APC^KO+N-APC^ exhibited reduced extrusion as compared to the WT and APC^KO^, and the reduction was again reverted by the addition of ATM1001. The data closely paralleled our observations from the laser-induced apoptosis, underscoring the suppression of cellular extrusion under conditions of elevated contractility and tissue tension (Figure 3d).

### APC N-terminus-induced increase in tension compromises apoptotic extrusion in intestinal organoid monolayers

Finally, we used mouse intestinal organoids to corroborate the results of our genome-edited cell lines. Small intestinal organoids were derived from APC^580^ mice, featuring loxP sites precisely integrated around APC exon 14 (Shibata et al. 1997). Exon 14 was then excised by *in vitro* Cre-recombination, resulting in the generation of truncated APC akin to the APC^Trunc^

MCF10A model utilised in our study (Figure 4a, S1a). Henceforth, we will refer to the APC-mutated organoids as APC^fl/fl^.

**Figure 4:**
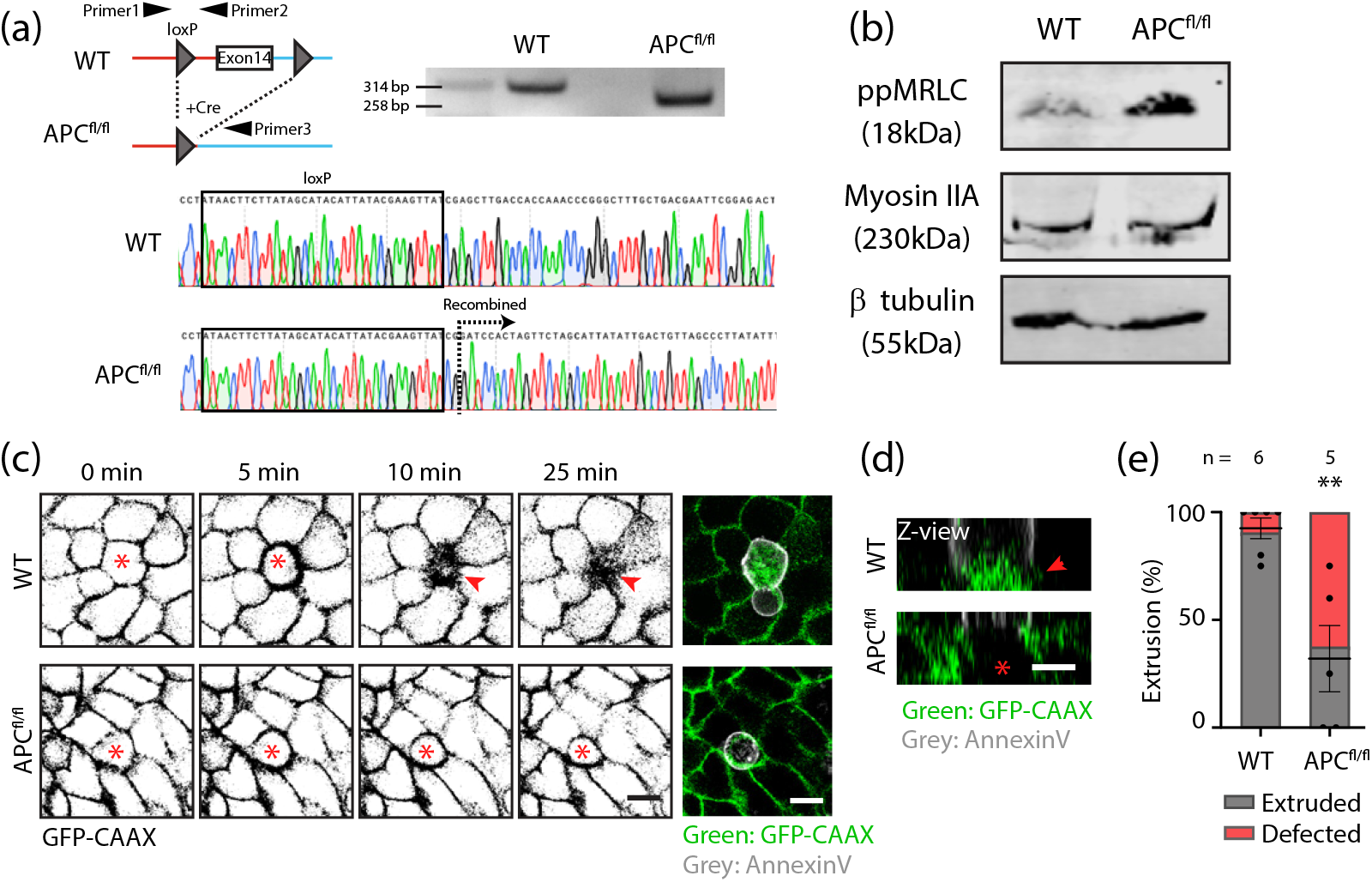
APC N-terminus-induced increase in tension compromises apoptotic extrusion in intestinal organoid monolayers. (a) Illustration of *in vitro* Cre recombination resulting in the deletion of *APC* exon 14, causing a genomic DNA PCR product shift from 314 to 258 bp. Sanger sequencing further confirmed recombination. (b) Western blot analysis of WT and APC^fl/fl^ organoid extracts detected for phosphorylated myosin light chain (ppMRLC), myosin IIA, and β tubulin. (c) Live cell recording of apoptotic cell extrusion within GFP-CAAX (green) expressing WT and APC^fl/fl^ organoid monolayers. Annexin V cell death marker (white). *cell injured with two-photon pulsed laser; Red arrow, closure of the epithelial layer upon extrusion. Scale bar: 10µm. (d) Orthogonal views of organoid monolayers in (c) at 25 min. ^*^retained cells after laser injury; Red arrow: closure of the epithelial layer upon extrusion. Scale bar: 10µm. (e) Quantification of apical cell extrusion as a result of laser injury as in (c) and (d). ^**^: p<0.01; Mann-Whitney U test. Data are means ± SEM, with individual data points indicated, and are obtained from 5-6 independent experiments.

Western blot analysis revealed that APC^fl/fl^ organoids demonstrated elevated ppMRLC compared to the WT counterparts (Figure 4b), indicating increased tissue contractility within the APC^fl/fl^ organoids, mirroring what we observed with the APC^Trunc^ MCF10A model. Then, we investigated whether cell extrusion is impaired within the APC^fl/fl^ organoids. To test this, we cultured organoids as 2-dimensional monolayers, using the protocol of Xi et al. (2022). Dissociated organoids were seeded onto cross-linked BME gels with a stiffness of approximately 400Pa, thereby facilitating optimal growth of intestinal organoids as 2D monolayers (Xi et al. 2022). Employing this method, we observed a reduction in cell extrusion upon laser injury to 37% within the APC^fl/fl^ monolayer, compared to 90% in the WT monolayer (Figure 4c-e, Movie 3). These findings support the data from MCF10A cells, confirming that cellular extrusion is impeded under conditions of APC N-terminus-induced elevation in tissue contractility. This further suggests that APC^Trunc^ has a capacity to inhibit apical extrusion by elevating tissue tension which can operate in different epithelial tissues, highlighting the generality of our findings.

Overall, our study suggests a model wherein a N-terminal APC truncation induces heightened tissue tension, resulting in morphological alterations and compromised apical cell extrusion within epithelial monolayers. Central to this mechanism is the critical role of RhoA in regulating epithelial tension by modulating the actomyosin cytoskeleton. Upon activation, RhoA triggers the downstream activation of ROCK, which enhances actomyosin contractility by phosphorylating MRLC and inhibiting Myosin Light Chain Phosphatase (MLCP) (Leerberg et al. 2014; Kimura et al. 1996). Our findings demonstrate that the expression of the N-terminal APC truncation mutant indeed amplifies tissue tension via the RhoA-ROCK pathway. Significantly, inhibition of cell contractility by the orthogonal strategy of inhibiting tropomyosin effectively restores tissue tension, reverses morphological changes and rescues apoptotic cell extrusion.

A key question for future research is to elucidate how expression of APC^Trunc^ stimulates RhoA signaling within cells. One possibility is that expression of this APC mutant activates a RhoA Guanine nucleotide exchange factor (GEF). The Armadillo repeats region within the N-terminus of APC has been reported to activate the GEF Asef (Kawasaki et al. 2000; Kawasaki et al. 2009). Asef, in turn, has been demonstrated to activate Rac1, CDC42 and also RhoA (Kawasaki et al. 2000; Gotthardt and Ahmadian 2007; Kim et al. 2018). Full-length APC undergoes a self-folding mechanism that auto-inhibits its N-terminus, thereby regulating Asef activation (Yang et al. 2021). However, the truncated N-terminus has been demonstrated to effectively activate Asef (Kawasaki et al. 2000). Understanding whether Asef or another RhoA GEF is activated by APC^Trunc^ is crucial for elucidating how this APC mutation impacts epithelial tissue mechanics and homeostasis. Our findings underscore the importance of considering APC-mediated regulation of tissue tension as a contributing factor to disease pathology.

## MATERIALS AND METHODS

### Cell culture

MCF10A was maintained in DMEM/F12 (Invitrogen; catalog #11330-032) containing 5% heat-inactivated horse serum (Invitrogen; catalog #16050-122), 20ng/ml human epidermal growth factor, 0.5mg/ml hydrocortisone, 100ng/ml cholera toxin, 10μg/ml insulin, and 100U/ml penicillin/streptomycin under standard cell culture condition (37°C, 95/5% air/CO_2_).

Small intestinal organoids were harvested from the APC^580s^ mouse as described before (Shibata et al. 1997; O’Rourke et al. 2016). The organoids were encapsulated in 50% Cultrex basement membrane extract type 2 (BME-II) (R&D Systems; catalog #3533-010-02) and maintained in advanced DMEM/F12 (Invitrogen; catalog #12634-010) supplemented with 50% medium conditioned with Wnt-3A, R-spondin 3 and noggin, 20% fetal bovine serum, and 100U/ml penicillin/streptomycin. 10μM of Y27632 (Watanabe et al. 2007) and 10μM of SB431542 (Miyoshi et al. 2012) were supplemented for the first 24 hours after passaging to increase viability, followed by changing to\ 2μM of IWP-2 and 2mM of valproic acid (Yin et al. 2014) and culture for three days for the enterocyte differentiation.

### APC deletion

APC knock-out (APC^KO^) and truncated (APC^Trunc^) MCF10A cells were generated through CRISPR/Cas9-mediated gene modification. sgRNAs targeting APC exon 1 5’-GCAAGTTGAGGCACTGAAGA-3’ and exon 11 5’-GCTTTGACAAACTTGACTTT-3’ (Saito-Diaz et al. 2018) were obtained from Integrated DNA Technologies. The sequences were cloned into pSpCas9(BB)-2A-Puro (PX459) vector (Addgene plasmid #62988), followed by lipofectamine-mediated transfection into the MCF10A cells. Cells were then selected in 1μg/ml puromycin for 48 hours and single clones were sorted through flow cytometry. Genotypes of the single clone were confirmed by PCR and sanger sequencing of *APC* exon 1 and exon 11 using the following primers: exon 1: 5’-CTTATAGGTCCAAGGGTAGCCAAG-3’ and 5’-TAAAAATGGATAAACTACAATTAAAAG-3’; exon 11: 5’-GATGATTGTCTTTTTCCTCTTGC-3’ and 5’-CTGAGCTATCTTAAGAAATACATG-3’.

APC^fl/fl^ small intestinal organoids were generated through Cre-loxP-mediated recombination by adding TAT-Cre recombinase *in vitro* to dispersed organoids for 8 hours, followed by encapsulating the organoids back into BME-II gel (Chan et al. 2023). The organoids were then cultured in non-conditioned medium for at least a week to select the clusters with successful recombination. Successful recombination was confirmed by gel electrophoresis and sanger sequencing of the the genomic DNA PCR product using the following primers: 5’-GTTCTGTATCATGGAAAGATAGGTGGTC-3’, 5’-CACTCAAAACGCTTTTGAGGGTTGATTC-3’, and 5’-GAGTACGGGGTCTCTGTCTCAGTGAA-3’ (Shibata et al. 1997).

### Viral transduction

Stable cell lines expressing E-cadherin-mCherry, APC^KO^ cells expressing N-APC-GFP (APC^KO+N-APC^), and organoids expressing eGFP-CAAX were generated through lentiviral transduction. Lentiviral construct pLL5.0 E-cadherin shRNA/mE-cadherin-mCherry (Addgene plasmid #101282), pLL5.0-eGFP-CAAX (Addgene plasmid #187252) or pLL5.0-N-APC-GFP, together with third-generation lentiviral packaging vectors (Addgene plasmids #12251, #12253, and #12259) were transfected into HEK293T cells to generate viral particles. Secreted lentivirus was collected at day 2 and 3 post transfection. After addition of polyethylene glycol 6000 (PEG 6000), virus was precipitated by centrifugation at 1500g for 1 hour at 4°C. Next, the virus was added to target cells or dissociated organoids and incubated 48 or 24 hours respectively, in the presence of 10μg/ml polybrene. mCherry-positive cells were selected through flow cytometry, whereas GFP-positive organoids were manually selected using a stereomicroscope.

### PDMS generation

Equal amount of Cy-A and Cy-B (Dow Corning) were mixed in a 50ml tube. Approximately 0.56 grams of the PDMS mixture was added onto a 35mm glass bottom dish, spin coat at 500 rpm for 30 seconds, and cured at 80°C for 2 hours. Cured PDMS was coated with 5% BME-II gel overnight at 4 °C, followed by incubation in 1% pluronic acid for 30 minutes. Dish was thoroughly washed with PBS before cell seeding.

### Western blotting

Cells grown to full confluency within a 6-well plate or organoids isolated from the BME-II gel were lysed using lysis buffer (50mM Tris-Cl, 2% sodium dodecyl sulfate (SDS), 0.1% bromophenol blue, 10% glycerol, 0.04M DTT, and 1xphosSTOP). After heating the lysate at 98°C for 10 minutes, the proteins were separated on polyacrylamide gels. The proteins were then transferred onto methanol-activated polyvinylidene difluoride (PVDF) membranes, followed by blocking with 5% skim milk for 1 hour at room temperature. Next, the membranes were incubated in primary antibodies (rabbit polyclonal antibody against human APC (cell signaling; catalog #2504), rabbit polyclonal antibody against ser19-phospho specific myosin light chain (Chemicon; catalog #AB3381), or rabbit polyclonal antibody against non-muscle myosin II heavy chain A (BioLegend; catalog #PRB-440P)) diluted in 5% bovine serum albumin (BSA) in TBS overnight at 4°C. The blots were then washed and probed with secondary antibodies (Goat anti-mouse IRDye 680RD or goat anti-rabbit IRDye 800CW) diluted in 5% skim milk for 1 hour at room temperature. After washing, the blots were visualised using a LI-COR Odyssey CLx system.

### Immunofluorescence staining

Cells grown to full confluency on coverslips were pre-permeabilized in cytoskeleton stabilisation buffer (10mM PIPES, 50mM NaCl, 3mM MgCl_2_, 300mM sucrose, 1x protease inhibitor) containing 0.5% triton X-100 for 5 minutes on ice, followed by fixation in 4% paraformaldehyde diluted in PBS at room temperature for 10 minutes. After washing 3 times with PBS, the fixed cells were blocked in 3% BSA for 1 hour at room temperature and incubated in primary antibodies (rat monoclonal antibody against α-catenin M-domain (α18, gifted from Dr A. Nagafuchi, Kumamoto University, Japan); mouse monoclonal antibody against α-catenin (Invitrogen; catalog #13-9700); Rabbit polyclonal antibody against α-catenin ABD (VD7, gifted from Professor Dr D. Vestweber, Max Planck Institute for Molecular Biomedicine, Germany); rabbit polyclonal antibody against vinculin (abcam; catalog #ab91459); rat monoclonal antibody against E-cadherin (Zymed; catalog #13-1900)) overnight at 4°C. Cells were washed 3 times with PBS, followed by incubation in secondary antibodies (goat antibodies conjugated with Alexa Fluor 488, 546 or 647 (Invitrogen)) for 1 hour at room temperature. Lastly, cells were washed 3 times with PBS and mounted onto glass slides with Prolong gold antifade mountant (Invitrogen; catalog #P36930). Imaging was performed using a Zeiss LSM880 confocal microscope.

### Laser injury

Cells were grown on 35mm glass bottom dishes to 90% confluency and imaged in Hank’s balanced salt solution (Sigma-Aldrich; catalog #H8264) supplemented with 5% FBS, 15mM HEPES, 5mM CaCl_2_, and 10mM glucose. For the tropomyosin inhibition, cells were incubated with 2.5µM ATM1001 for 24 hours and treatment was maintained throughout imaging.

Organoids were harvested from BME-II gel by adding cold trypLE solution (Gibco; catalog #12604013) to the gels. The organoids were further dissociated in the solution by incubating for 3 minutes at 37 °C, followed by pipetting up and down 30 times. The dissociated organoids were then seeded onto BME-II gel cross-linked with N-hydroxysuccinimide (NHS) and 1-Ethyl-3-(3-dimethylaminopropyl)carbodiimide (EDC) (Xi et al. 2022), and cultured as 2D monolayers until confluent.

Imaging cells and organoid monolayers was performed on a Zeiss LSM710 confocal microscope equipped with a MaiTai eHP tuneable 760-1040nm laser. The nuclei of the cells were ablated with a 790nm laser (70% laser power, 30 iteration) and the process of extrusion was recorded for 1 hour at an interval of 30 seconds. Apoptosis of the injured cells was confirmed using Annexin V conjugated with Alexa Fluor 647 (Thermo Fisher Scientific; catalog #A23204).

### Etoposide assay

Cells grown to full confluency were incubated with 500µM etoposide (Adooq Biosciences LLC #A10373) for 6 hours at 37°C. Next, cells were fixed with 4% paraformaldehyde diluted in PBS at room temperature for 10 minutes and permeabilised with 0.1% triton X-100 for 10 minutes, followed by immunofluorescence staining. Caspase 3 (Cell signaling; catalog #9661) was used to identify the apoptotic cells and E-cadherin mAb antibody (Zymed; catalog #13-1900) was used to highlight the borders of individual cells. Imaging was performed on a Zeiss Axio Observer 7 microscope.

### GTP-RhoA pulldown

Pulldown of GTP-RhoA was performed according to the manufacturer’s instruction (Cytoskeleton; catalog #BK030). In brief, cells were serum-starved for 24 hours, followed by serum stimulation for 30 mins. Next, cells were washed with ice-cold PBS, lysed, and incubated with 15µl Rhotekin-RBD affinity beads for 2 hours at 4 °C with rotation. The beads were then collected and washed. Bound GTP-RhoA was extracted by heating the beads in western blot lysis buffer for 5 mins and the lysates were analysed by western blot.

### Image and statistical analysis

Images were analysed using FIJI (ImageJ). Immunofluorescent images were Z-projected as the sum of intensity. Junctional marker (α-catenin or E-cadherin) signals were used to create a mask after a gaussian blur was applied. The mask was projected onto each channel to obtain average intensities of the various stainings.

Ratiometric measurements were obtained by dividing the average intensities of the tension-sensitive protein by the average intensities of the total junctional protein. Ratios were further normalised against the WT control.

For laser injury, cell was classified as extruded if the neighbouring cells successfully closed the gap underneath the annexin V-positive injured cell within 1 hour of imaging. Otherwise, the extrusion was classified as non completed.

All numerical data are presented as mean ± standard error of mean. Statistical analysis was performed using GraphPad Prism. Statistical parameters for individual experiments are listed in the corresponding figure legends. This includes the statistical test performed, sample size (n), number of independent experiments, and statistical significance.

## AUTHOR CONTRIBUTIONS

The study is conceptualised by W.J.G, A.S.Y, and I.N. W.J.G performed all the experiments and analysis under the supervision of A.S.Y and I.N. R.G, J.B, and H.E.A provided advice for the works related to organoids. E.C.H and P.G provided the tropomyosin inhibitor ATM1001 used in the study.

## ACKNOWLEDGEMENTS

Our work was supported by grants from the National Health and Medical Research Council of Australia (GNT2010704) and the Australian Research Council (DP220103951 and ARC Laureate Fellowship FL2023000092) to A.S.Y; grants from the National Health and Medical Research Council of Australia (1079866 and 1100202) and the Australian Department of Industry, Science and Resources (CRC-P-355) to P.W.G and E.C.H; a grant from the National Health and Medical Research Council of Australia (2021181) to H.E.A; a fellowship from the European Molecular Biology Organization (EMBO <u>ALTF 251-2018</u>) to I.N. Microscopy was performed at the ACRF/IMB Cancer Research Imaging Facility created with the generous support of the Australian Cancer Research Foundation.

## DISCLOSURE OF POTENTIAL CONFLICTS OF INTEREST

E.C. Hardeman reports receiving a commercial research grant from and has ownership interest (including patents) in TroBio Therapeutics Pty Ltd. P.W. Gunning reports receiving other commercial research support from and has ownership interest (including patents) in TroBio Therapeutics Pty Ltd.

## FIGURE AND MOVIE LEGENDS

<u>Supplementary figure 1.</u>

(a) Illustration of APC full length protein (2843 amino acids), APC^Trunc^ (511 amino acids), N-APC (641 amino acids) tagged with eGFP, and APC^fl/fl^ (580 amino acids).

(b) Western blot of WT, APC^KO^, and APC^Trunc^ cell extracts detected for full-length APC and β tubulin.

(c) Western blot analysis and quantification of WT and APC^Trunc^ cell extracts detected for pulldown RhoA-GTP, total RhoA, and β tubulin. N=2 independent experiments; ^*^P=0.0298; student’s unpaired t-test. Data are means ± SEM, with individual data points indicated.

(d) Western blot of WT and APC^KO+N-APC^ cell extracts detected for GFP and β tubulin.

<u>Movie 1.</u>

Phase contrast live cell imaging of (a) WT, (b) APC^Trunc^, (c) Y27632-treated APC^Trunc^, and (d) Y27632 wash off APC^Trunc^ MCF10A cells related to figure 1. Scale bar: 50µm.

<u>Movie 2.</u>

Laser-induced apoptotic cell extrusion of E-cadherin-mCherry (red) expressing (a) WT, (b) APC^KO^, (c) APC^Trunc^, (d) APC^KO+N-APC^, and (e) ATM1001-treated APC^Trunc^ MCF10A cells related to figure 3. Green: N-APC-GFP expression; White: annexin V cell death marker. Scale bar: 10µm.

<u>Movie 3.</u>

Laser-induced apoptotic cell extrusion of GFP-CAAX (green) expressing (a) WT, and (b) APC^fl/fl^ organoid monolayers related to figure 4. White: annexin V cell death marker. Scale bar: 10µm.

