## Supplementary figures and images for "A truncation mutant of adenomatous polyposis coli (APC) impairs apical cell extrusion through elevated epithelial tissue tension"

### Supplemental figure

Supplementary figure 1:

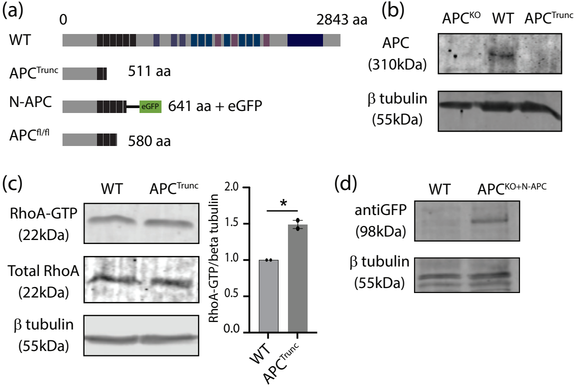
